# SAMBA controls the rate of cell division in maize development through APC/C interaction

**DOI:** 10.1101/2021.04.22.440954

**Authors:** Pan Gong, Michiel Bontinck, Kirin Demuynck, Jolien De Block, Kris Gevaert, Dominique Eeckhout, Geert Persiau, Stijn Aesaert, Griet Coussens, Mieke Van Lijsebettens, Laurens Pauwels, Geert De Jaeger, Dirk Inzé, Hilde Nelissen

## Abstract

SAMBA has been identified as a plant-specific regulator of the anaphase-promoting complex (APC/C) which controls unidirectional cell cycle progression in Arabidopsis, but so far its role was not studied in monocots. Here, the association of SAMBA with APC/C was shown to be conserved in maize. Two *samba* CRISPR alleles showed growth defects that aggravated with plant age such as dwarfed plants due to shortened upper leaf length, erect leaf architecture, and reduced leaf size due to an altered cell division rate and cell expansion. Despite the fact that in both alleles the frameshift occurred at the same position, the two alleles differed in the severity and developmental onset of the phenotypes, because *samba-1* represented a knock-out allele, while translation re-initiation in *samba-3* resulted in a truncated protein that was still able to interact with the APC/C and regulate its function, albeit with altered APC/C activity or efficiency. Our data are consistent with a dosage-dependent role for *SAMBA* to control developmental processes for which a change in growth rate is pivotal.

## Introduction

The shoot apical meristem (SAM) gives rise to all above-ground plant organs. Leaves initiate when leaf primordia emerge from the flanks of the SAM. A typical characteristic of monocot leaf development is the sheathing leaf base leading to older leaves to be wrapped around younger developing leaves (Conklin et al., 2019). The leaf sheath is separated from the leaf blade by the auricle and the ligule. Auricles are two wedge-shaped structures on either side of the mid-vein, which act as hinges to allow the leaf blade to project at an angle from the vertical leaf sheath axis. The size of the auricles and ligule has been shown to be a major determinant of the leaf angle (Kong et al., 2017). When the leaf becomes visible from the whorl, two key growth parameters, leaf elongation rate (LER) or maximal growth rate and leaf elongation duration (LED) or how long maximal growth is maintained can be determined by daily measuring its length until it is maximal (Nelissen et al., 2013). The leaf is attached at the node which is a structure that is derived from the intercalary meristems that also give rise to the internodes, which determine the final plant height (Tsuda et al., 2017).

Both leaves and internodes grow post-embryonically through cell division and cell expansion. The contribution of cell division to the final organ size is determined by the number of dividing cells and the cell division duration and rate that is controlled by the cell cycle progression. The cell cycle is one of the most highly conserved processes among eukaryotes and is a strictly controlled process in which a cell duplicates its genome and equally distributes it between its two daughter cells. To ensure correct duplication and distribution of the genome and separation of the two daughter cells, the cell cycle has several checkpoints, which only allow progression through the cell cycle if the previous step has been correctly completed (De Veylder et al., 2007). Ubiquitin-mediated degradation of essential checkpoint proteins is a major mechanism ensuring unidirectional progression through the cell cycle checkpoints (Eloy et al., 2015; Sharma et al., 2016), for which E3 ligases provide substrate specificity. The SKP/CUL/RBX/F-box (SCF) protein complex and the anaphase-promoting complex/cyclosome (APC/C) play a major role in the regulation of cell cycle progression by ubiquitin-mediated degradation (Eloy et al., 2015). The APC/C is a large multi-subunit complex that is conserved between yeast, mammals and plants (Lima et al., 2010; Eloy et al., 2015) and consists of a catalytic and substrate recognition module, a tetratricopeptide repeat (TPR) lobe and a platform module. The catalytic core of the APC/C is comprised of APC2 and APC11(Gmachl et al., 2000; Leverson et al., 2000; Tang et al., 2001). Substrate specificity of the catalytic core is obtained through association with APC10 and the CELL DIVISION CYCLE 20 (CDC20) and CELL CYCLE SWITCH 52 (CCS52) families of APC/C co-activator proteins (Da Fonseca et al., 2011; Chang et al., 2015).

During the mitotic cell cycle, the APC/C has been shown to target cell cycle regulators such as CYCLINs (CYCs), CYCLIN-DEPENDENT KINASE inhibitors, and proteins involved in DNA replication (Castro et al., 2005). The ability of the APC/C to mark proteins for degradation depends on the presence of destruction signals (degrons), such as the D-box or KEN-box motifs, in its target proteins (Glotzer et al., 1991; Pfleger and Kirschner, 2000). Besides mitotic cell division, the APC/C is also involved in regulating endoreduplication, which is a different type of cell cycle consisting of several rounds of DNA synthesis without subsequent cell division (De Veylder et al., 2011). The onset of endoreduplication often coincides with the exit from the mitotic cell cycle and the onset of cell differentiation (Breuer et al., 2010). Furthermore, the endocycle may serve as a mechanism to sustain cellular expansion without cell division, as endocycle rates often correlate with final cell size (Melaragno et al., 1993), although this is not always the case (Beemster et al., 2002; De Veylder et al., 2011). Furthermore, several plant-specific APC/C inhibitors have also been identified, such as SAMBA (Eloy et al., 2012). In Arabidopsis, SAMBA has been shown to directly interact with APC3b and the *SAMBA* gene was highly expressed during early development (Eloy et al., 2012). *SAMBA* loss of function results in increased cell proliferation, leading to larger meristem size as well as the formation of larger leaves, roots and seeds, by shortening the cell cycle duration (Eloy et al., 2012).

The CRISPR/CAS9 system allows to efficiently introduce double-strand DNA breaks (DSBs) to a specific target site in the genome. Non-homologous end-joining (NHEJ) is the main mechanism to repair these DSBs, which can introduce small insertions or deletions (indels), resulting in a shift in the open reading frame, leading to a premature stop codon (PTC) in the expressed transcript. Normally, the aberrant mRNAs are degraded by nonsense-mediated decay (NMD) resulting in a loss-of-function (Doudna and Charpentier, 2014). Recently, multiple studies further analyzed the post-transcriptional effect of indels on the target gene, showing that alternative splicing (skipping the mutated exon) and/or translation re-initiation downstream of the CRISPR frameshift mutation can produce truncated functional proteins in cell lines, animals and plants (Tang et al., 2018; Smits et al., 2019). In approximately one third of 136 CRISPR target genes in HAP1 cells residual proteins were found due to skipping of the mutated exon or translation re-initiation (Smits et al., 2019). Also in rice, *de novo* alternative splicing of *Auxin/Indole-3-Acetic Acid 23* (*OsIAA23*) circumvented the premature stop and thereby preserved the wild-type phenotype in a CRISPR/Cas9 mutant (Jiang et al., 2019).

Here, we found that the interaction of SAMBA and APC/C was conserved in maize and remained stable in the cell division and cell expansion zone. CRISPR/Cas9 mutants in the maize *SAMBA* ortholog displayed dwarfism, erect upper leaves, reduced organ and tissue growth, but the onset and severity of the phenotypes was altered in two alleles although the frameshift occurred at the same position. The difference in the allelic strength was caused by the formation of a truncated protein due to translation re-initiation in one allele, while no detectable protein or a very short N-terminal peptide was formed in the null allele. Phenotypic analysis showed that the knock-out *SAMBA* mutant had a higher cell production by increased cell division rate but reduced cell size, and mainly affected developmental processes for which a change in growth rate is instrumental, such as ligule formation and internode elongation. The truncated SAMBA protein retained partial functionality and was still able to interact with the APC/C. In addition, the partial complementation of the null allele by the truncated protein showed that SAMBA functioned in a dose-dependent manner to control the cell division rate.

## Results

### SAMBA interacts with the APC/C complex *in vivo*

To verify whether the maize SAMBA ortholog also interacts with the APC/C, the full-length *SAMBA* coding sequence was fused to the GS^rhino^ affinity tag (Nelissen et al., 2015) (short as *SAMBA-GS*) and ectopically expressed into the maize inbred line B104. Affinity purification followed by mass spectrometry (AP/MS) was performed on the embryogenic callus, identifying orthologs of the core components of the APC/C which were previously identified as interacting proteins with SAMBA in Arabidopsis (Eloy et al., 2012) (Table 1, Supplemental dataset 1). Furthermore, APC15b, previously not identified in Arabidopsis was enriched in maize callus. In addition to these known components and regulators of the APC/C, 56 proteins that were significantly enriched compared to the control purifications were identified. None of these proteins were previously found to interact with SAMBA in Arabidopsis, but some were already to be reported as putative targets of APC/C in human cells (Horn et al., 2011) (Table S1, Supplemental dataset 1). Among these proteins are several members of the DYNAMIN-RELATED PROTEIN (DRP) family, DRP1A, DRP1E, DRP2A, DRP2B and DRP5B. Furthermore, CALLOSE SYNTHASE, which is involved in cell plate synthesis and is regulated by DRP proteins (Verma, 2001) was retrieved as SAMBA interacting protein.

**Table 1:**
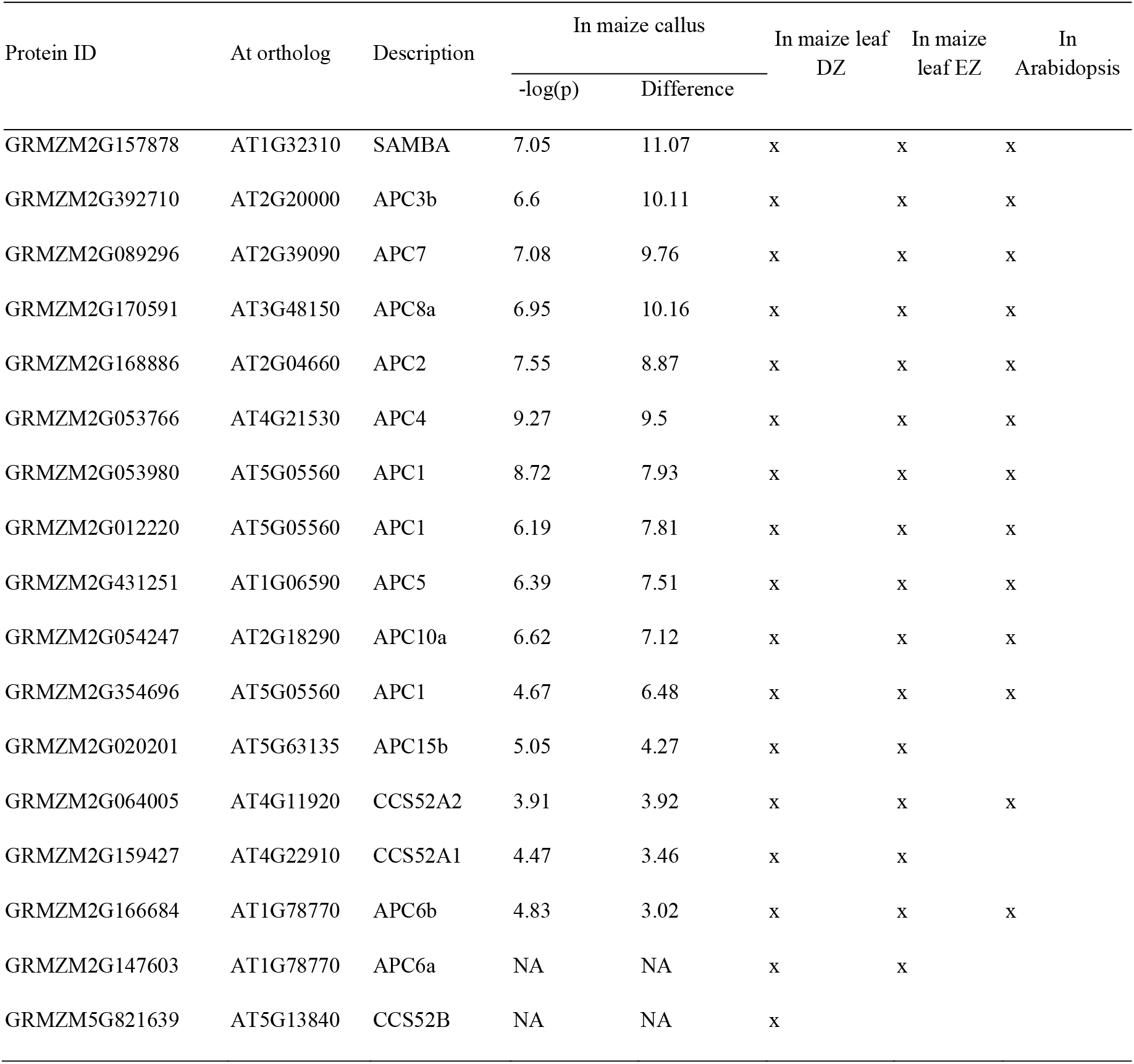
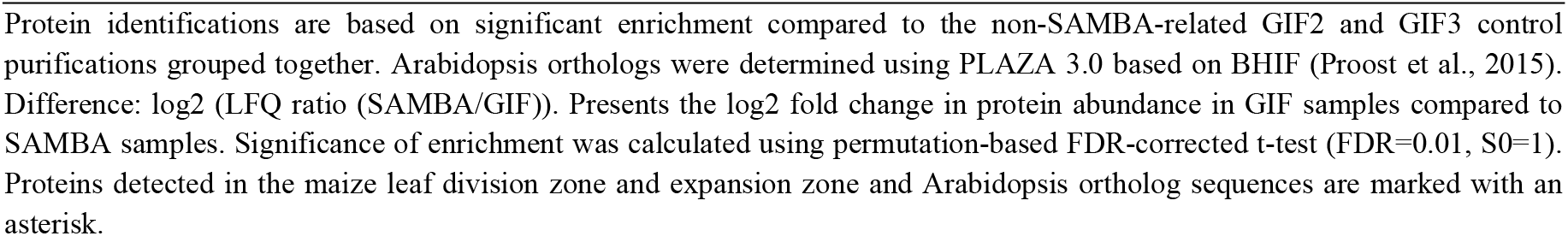
Known APC components identified through AP/MS from maize callus, leaf division zone (DZ) and expansion zone (EZ) using SAMBA-GSrhino as bait

| Protein ID | At ortholog | Description | In maize callus |  | In maize leaf<br>DZ | In maize<br>leaf EZ | In<br>Arabidopsis |
| --- | --- | --- | --- | --- | --- | --- | --- |
|  |  |  | -log(p) | Difference |  |  |  |
| GRMZM2G157878 | AT1G32310 | SAMBA | 7.05 | 11.07 | x | x | x |
| GRMZM2G392710 | AT2G20000 | APC3b | 6.6 | 10.11 | x | x | x |
| GRMZM2G089296 | AT2G39090 | APC7 | 7.08 | 9.76 | x | x | x |
| GRMZM2G170591 | AT3G48150 | APC8a | 6.95 | 10.16 | x | x | x |
| GRMZM2G168886 | AT2G04660 | APC2 | 7.55 | 8.87 | x | x | x |
| GRMZM2G053766 | AT4G21530 | APC4 | 9.27 | 9.5 | x | x | x |
| GRMZM2G053980 | AT5G05560 | APC1 | 8.72 | 7.93 | x | x | x |
| GRMZM2G012220 | AT5G05560 | APC1 | 6.19 | 7.81 | x | x | x |
| GRMZM2G431251 | AT1G06590 | APC5 | 6.39 | 7.51 | x | x | x |
| GRMZM2G054247 | AT2G18290 | APC10a | 6.62 | 7.12 | x | x | x |
| GRMZM2G354696 | AT5G05560 | APC1 | 4.67 | 6.48 | x | x | x |
| GRMZM2G020201 | AT5G63135 | APC15b | 5.05 | 4.27 | x | x |  |
| GRMZM2G064005 | AT4G11920 | CCS52A2 | 3.91 | 3.92 | x | x | x |
| GRMZM2G159427 | AT4G22910 | CCS52A1 | 4.47 | 3.46 | x | x |  |
| GRMZM2G166684 | AT1G78770 | APC6b | 4.83 | 3.02 | x | x | x |
| GRMZM2G147603 | AT1G78770 | APC6a | NA | NA | x | x |  |
| GRMZM5G821639 | AT5G13840 | CCS52B | NA | NA | x |  |  |

Protein identifications are based on significant enrichment compared to the non-SAMBA-related GIF2 and GIF3 control purifications grouped together. Arabidopsis orthologs were determined using PLAZA 3.0 based on BHIF (Proost et al., 2015). Difference: log2 (LFQ ratio (SAMBA/GIF)). Presents the log2 fold change in protein abundance in GIF samples compared to SAMBA samples. Significance of enrichment was calculated using permutation-based FDR-corrected t-test (FDR=0.01, S0=1). Proteins detected in the maize leaf division zone and expansion zone and Arabidopsis ortholog sequences are marked with an asterisk.

In addition, seven independent stable transgenic plants were obtained of which *SAMBA-GS4, SAMBA-GS5* and *SAMBA-GS6* showed a high accumulation of the fusion protein (Fig. S1). Using AP-MS, we mapped the dynamics of the SAMBA-interacting proteins in the division and expansion zone of growing *SAMBA-GS5* leaf four. All APC/C components identified in the maize callus, except APC6a and CCS52B were retrieved in the leaf tissue (Table 1). Label-free quantification of co-purified proteins showed that the abundance of SAMBA and of the majority of the identified APC/C subunits did not change significantly between both zones (Fig. S2). However, APC7 and even more pronounced CCS52B were found to be significantly enriched in the division zone (Fig. S2; Supplemental dataset 2). Furthermore, 20 out of the 56 additional proteins, which were identified after affinity enrichment from callus, were also identified after affinity enrichment from leaf tissue, including DRP1A, DRP1E, DRP2A, DRP2B, and DRP5B (Table S1). Nevertheless, none of these proteins were significantly enriched in either developmental subzone (Supplemental dataset 2).

### *samba* frameshift mutants are dwarfed with decreased organ size

Because ectopic expression of the *SAMBA-GS* in maize did not induce observable phenotypes on leaf and plant growth (Fig S1B-C), the function of *SAMBA* in maize development was investigated by generating mutants using CRISPR/Cas9 with a dual gRNA approach (Fig. 1A). Both gRNAs had a very high MIT Specificity Score (91% for the first gRNA and 96% for the second gRNA) in CRISPOR (Concordet and Haeussler, 2018) indicating a very low risk of off-targets. The *samba-1* allele contained a single nucleotide deletion at both target sites, which caused a frameshift in the *SAMBA* coding sequence, resulting in 22 missense amino acids (AAs) prior to a premature stop codon (Fig. 1B-C). The *samba-2* allele contained a 30-nucleotide deletion at the first target site, which caused an in-frame deletion of 10 AAs while no mutation occurred at gRNA2 (Fig. 1B-C). The *samba-3* allele contained a single T insertion at both target sites (Fig. 1B), resulting in a frameshift at the same location as the *samba-1* allele but with only three missense AAs prior to the premature stop codon (Fig. 1C). The T0 *samba* mutants were backcrossed with B104 and the T1 heterozygote plants without Cas9 were self-pollinated to obtain the T2 generation. The T3 plants obtained by self-pollinating of the T2 heterozygotes were further analysed. We characterized the phenotypes of the *samba* T3 plants 70 days after sowing by comparing them to wild type (WT) segregating siblings. The phenotype of *samba-2* was similar to the WT (Fig. 2A), while *samba-1* mutants were severely dwarfed (plant height was reduced with 80% compared to its WT (Fig. 2A-B), which was due to significantly decreased elongation from internode 9 onwards (Fig. 3A). The average mature cell length of *samba-1* internode 9 was significantly decreased by 70.8% (Fig. S3A). The *samba-3* plants displayed a semi-dwarf phenotype (plant height was decreased by 65% compared to its WT) because the internodes from internode 12 onwards were significantly reduced (Fig. 2A-B and 3A). The average mature cell length of *samba-3* internode 12 was significantly decreased by 28.9% (Fig. S3B). Both *samba-1* and *samba-3* plants transitioned from vegetative development into reproductive development but failed to produce seeds in reciprocal crosses with WT. Leaf development was also affected in the *samba* mutants. The sheath length of lower and higher leaves was significantly increased and significantly decreased, respectively, in *samba-1*, whereas all leaf sheath lengths were significantly increased in *samba-3* (Fig. 3B). Furthermore, both blade length and width were significantly reduced in *samba-1* and *samba-3*; for the leaf width this was significant (P< 0.05) earlier in *samba-1* than in *samba-3* (Fig. 3C-D). The average mature leaf 12 cell length were significantly reduced by 31.3% and 14.5% in *samba-1* and *samba-3*, respectively, compared to their WT (Fig. S3C-D). Also, the transition between the leaf sheath and lamina was affected in the *samba-1* and *samba-3* alleles. The ligule was elongated and elliptical in shape and the auricle size was extremely decreased in both mutants, resulting in an erect leaf architecture (Fig. 2C). The *samba-1* leaves showed this ligule phenotype from leaf 8 onward, whereas the *samba-3* ligule phenotype was visible from leaf 10 onward (Fig. 2C). Furthermore, the *samba* mutants showed a decreased root density, but similar root length compared to the WT (Fig. S4). To conclude, the *samba-1* and *samba-3* mutants showed similar growth phenotypes, but the onset of the phenotypes differed between the two alleles. The mutant phenotypes could be complemented to WT upon introgression of a *SAMBA* overexpression line, *SAMBA-GS5*, into both *samba* lines (Fig. 4), which confirmed that the growth defect was caused by the *samba-1* and *samba-3* mutations.

**Figure 1.**
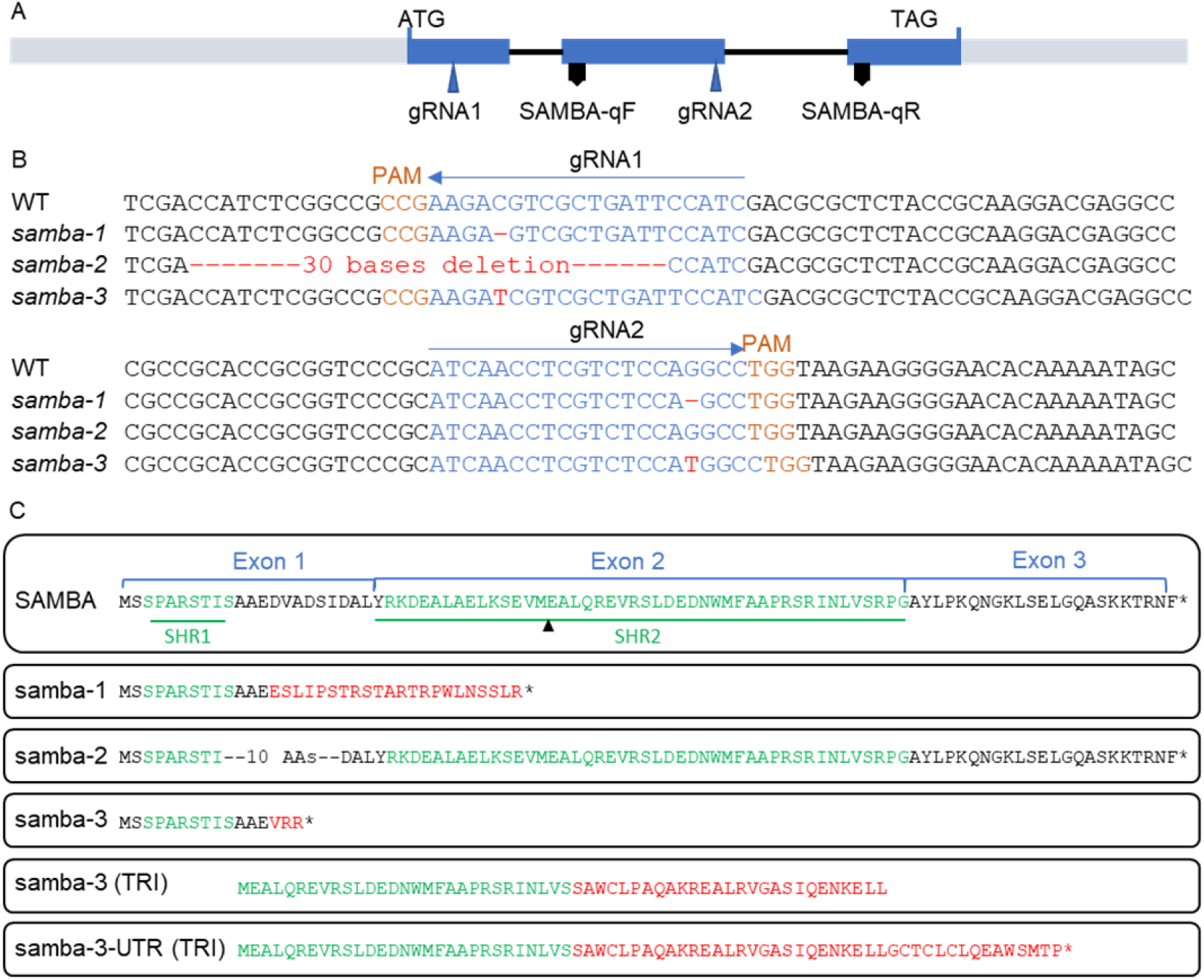
*samba* mutants obtained through CRISPR/Cas9 gene editing. **(A)** Structural representation of the maize SAMBA gene structure showing the target sites of the two gRNAs. **(B)** The gRNA sequences (blue), PAM (orange) and mutation sites (red) of the *samba* alleles. **(C)** The amino acid sequences of SAMBA WT and putative samba mutant isoforms. SAMBA homology region 1 (SHR1) and SHR2, as defined by Eloy (Eloy et al., 2012) are marked in green and the missense AAs are indicated red. SAMBA-qF and SAMBA-qR indicate the position of the primers used for SAMBA qRT-PCR. * represents a stop codon.

**Figure 2.**
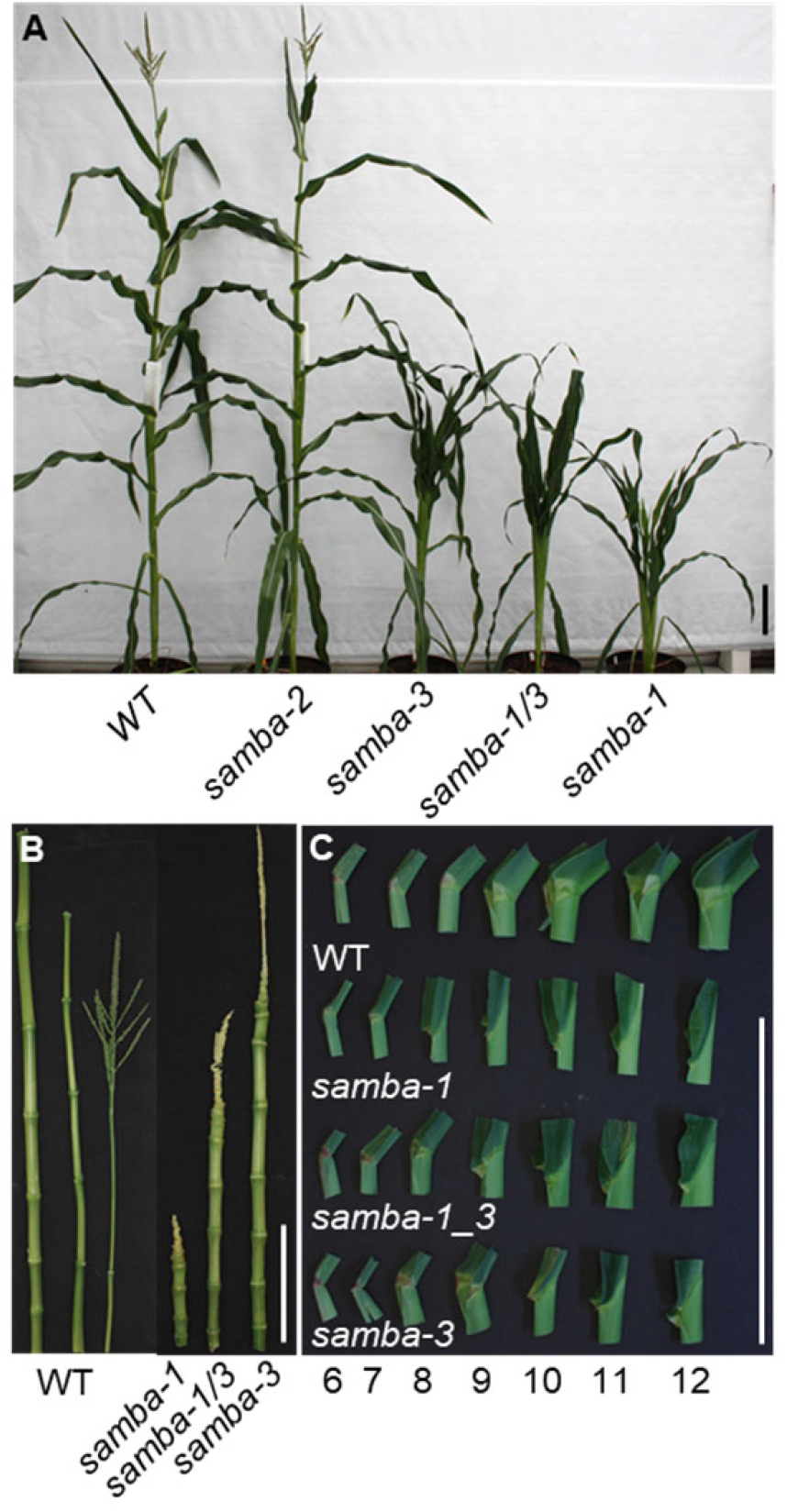
Overview of the phenotypes of WT and *samba* mutant plants. **(A)** WT and samba mutant plants grown in the greenhouse 70 days after sowing. The internodes **(B)** and ligule morphology **(C)** of WT plant and *samba* mutants from leaf 6 to leaf 12. Scale bar = 20 cm.

**Figure 3.**
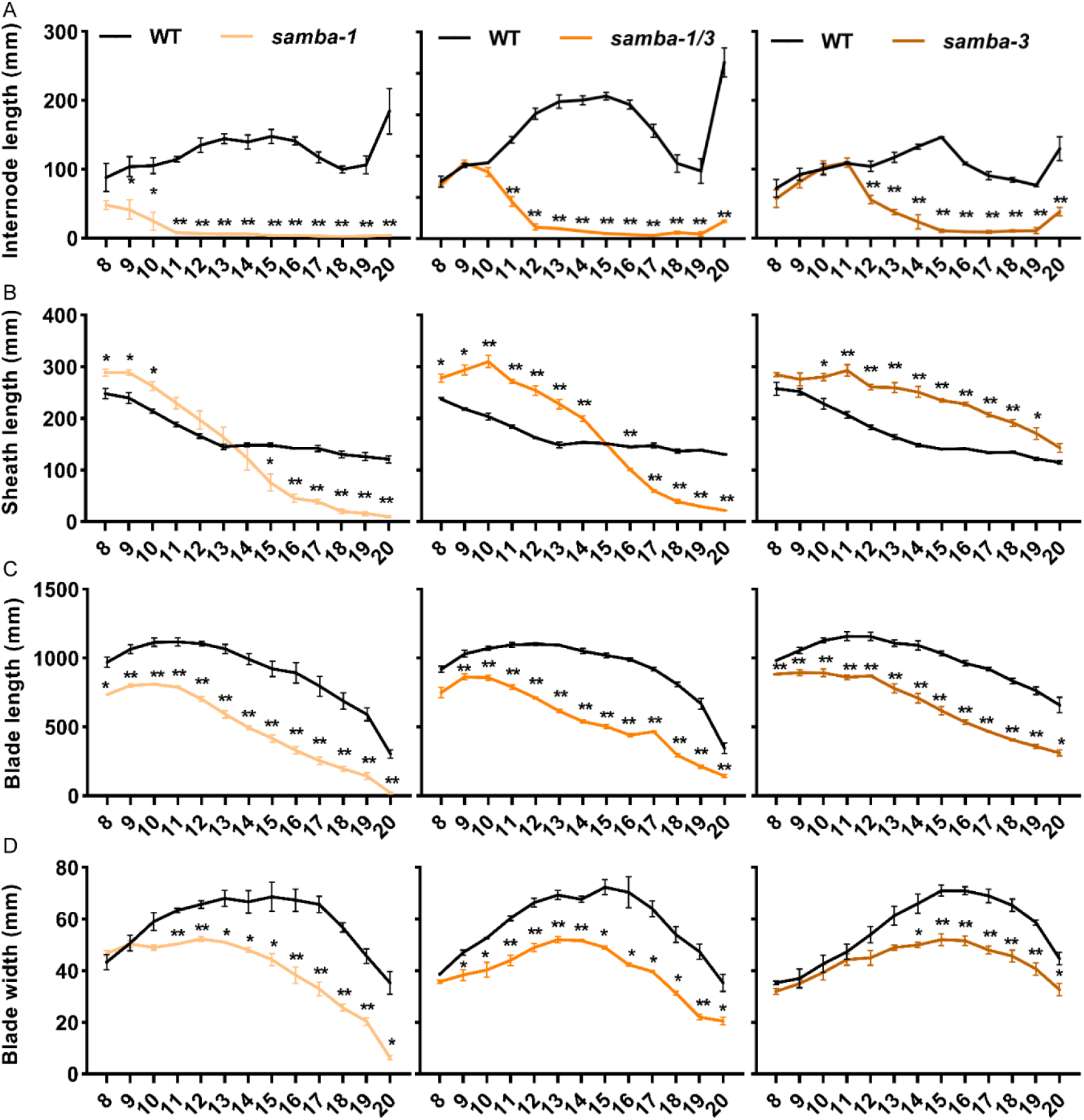
Quantification of internode length (A), leaf sheath (B) and blade (C) length and blade width (D) of the *samba* mutants in comparison to WT. Error bars represent standard error and significant differences were determined using Student’s *t*-test: *, P<0.05, **, P<0.01 (n≥3).

**Figure 4.**
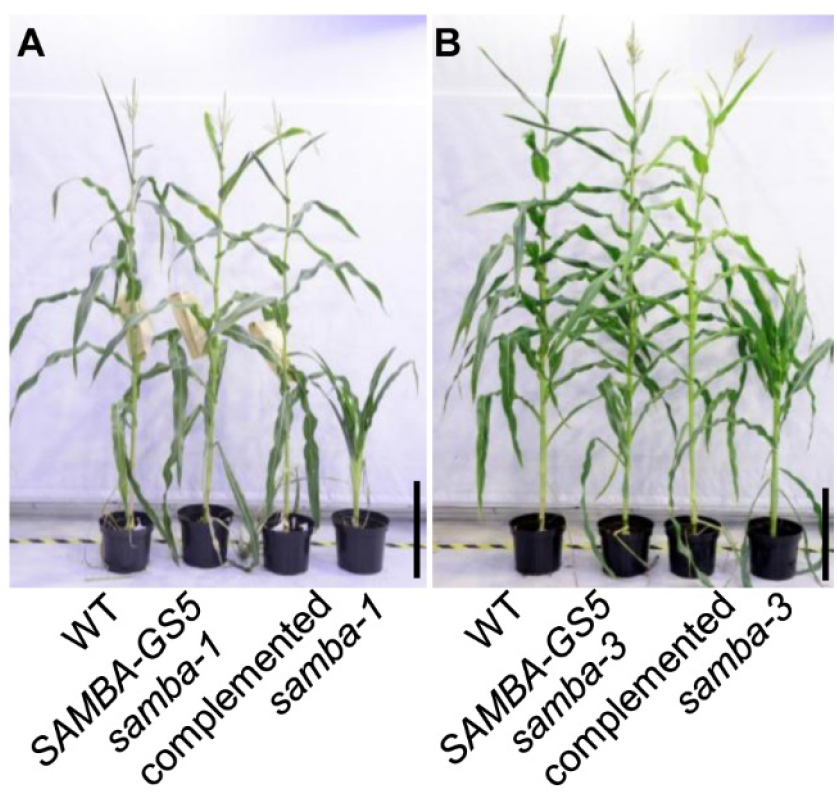
Complementation of the *samba* mutant plants by introducing *SAMBA-GS5*. Plants phenotype of WT, *SAMBA-GS5, samba* complemented plants and *samba-1* **(A)**, or *samba-3* **(B)**. Bars = 50cm.

Both *samba* alleles appeared to have a different strength and to analyze if their different onset of the phenotypes was due to a dosage effect, the heterozygous *samba-1* and *samba-3* mutants were crossed to generate the *samba-1/3* bi-allelic mutant. The bi-allelic *samba-1/3* mutant showed the significantly shortened internode length above internode 11, and the internode above the 15^th^ internode was barely elongated (Fig. 3A). Moreover, the elongated ligule was present at leaf 9, while the elliptical ligule was visible on leaf 8 and leaf 10 for *samba-1* and *samba-3*, respectively (Fig. 2C). These data show that the *samba-1/3* mutant showed the typic *samba* growth defects such as reduced plant height, shortened upper internode length, increased leaf sheath length, smaller and erect leaves and fewer roots compared to the WT, but the onset of the phenotypes was intermediate to that of the single mutants (Fig. 2-3; Fig. S4).

### *samba* mutants have an accelerated cell cycle and decreased cell size

To investigate the cellular effect of the *samba-1* mutation, the epidermal layer of internodes and mature leaves was imaged. Both WT and *samba-1* internode and leaf epidermal cells were organized in cell files parallel with the vertical internode axis or leaf mid-vein, but the cells of *samba-1* were smaller than those of the WT (Fig. S5A-B). Stomata developed normally in the *samba-1* mutant (Fig. S5C-D), but leaf blades showed altered leaf hair development (Fig. S5F). WT leaves developed long macrohairs and bicellular microhairs, spaced by prickle hairs (Fig. S5E, G), whereas in *samba-1* mutants, the large macrohairs were completely absent and a large number of enlarged prickle hairs were regularly spaced alongside the files of bulliform cells (Fig. S5F, H). The morphology and size of the *samba-1* leaf vascular bundles were similar to those of WT, but the number of minor veins per major vein was significantly reduced by 41% (Fig. S5I, J).

To gain more insights into defects in cell division or cell expansion, a kinematic analysis of the growing leaf four was performed in the most severe *samba-1* allele and the WT (Table 2). The kinematic analysis was based on daily leaf growth measurements, a DAPI staining to determine the size of the division zone and cell length profiles along the growth zone to quantify the contributions of cell division and/or cell expansion to the growth phenotype^5^. The fourth leaf of *samba-1* showed a moderate, but significant decrease of 6.64% (P<0.05) in final leaf length, while the LER was not significantly affected (Table 2). The effect of the *samba-1* mutation was much more pronounced at the cellular level, where a significant increase in cell production of 13.61% (P<0.05) was observed, while the mature cell size was reduced with 20.92% (P<0.05). This opposite effect on cell division and cell expansion explained the rather mild effect on final leaf length. The increase in cell production was caused by a significantly increased cell division rate (25.78%, P<0.05) and a decreased cell cycle duration (35.42%, P<0.05). The average cell cycle duration in WT leaves was 23.55 h while it was reduced to 17.4 h in the *samba-1* mutant leaves (Table 2). Flow cytometry on mature leaf 12 showed a slightly increased fraction of 2C cells relative to the population of 4C and 8C cells (Fig. S6A-B). This shift towards more 2C cells was more pronounced in developing internodes (Fig. S6C-D).

**Table 2:** Effect of *samba-1* mutation on cellular parameters of leaf 4 growth

| Parameter | WT | *samba-1* | P-value | % |
| --- | --- | --- | --- | --- |
| Final leaf length (mm) | 558.43 ± 5.80 | 523.67 ± 3.91 | 2.38E-05 | -6.64 |
| Mature cell size (µm) | 119.96 ± 1.15 | 99.21 ± 1.61 | 7.75E-04 | -20.92 |
| Cell production (cell h<sup>-1</sup>) | 17.40 ± 0.18 | 20.14 ± 0.38 | 0.01 | 13.61 |
| Cell division rate (cells cell<sup>-1</sup> h<sup>-1</sup>) | 0.030 ± 0.002 | 0.040 ± 0.002 | 0.03 | 25.78 |
| Cell cycle duration (h) | 23.55 ± 1.62 | 17.40 ± 0.89 | 0.04 | -35.42 |
| Division zone size (cm) | 1.49 ± 0.10 | 1.24 ± 0.06 | 0.11 | -19.84 |
| Expansion zone size (cm) | 2.88 ± 0.32 | 1.87 ± 0.08 | 0.08 | -53.83 |
| Average size of dividing cells (µm) | 26.91 ± 0.46 | 26.37 ± 0.34 | 0.4 | -2.06 |
| Leaf elongation rate (mm*h<sup>-1</sup>) | 2.08 ± 0.04 | 2.04 ± 0.04 | 0.19 | -2.31 |

Statistics were calculated based on Student's *t*-test, n=3.

### The frameshift mutation in *samba-3* leads to a partially functional truncated protein by translation re-initiation

To understand the differences in the severity and onset of the growth phenotypes between the different *samba* alleles, we analyzed the *SAMBA* mRNA level by qRT-PCR. In *samba-1*, the relative expression of *SAMBA* was reduced by 83% and 74% in leaf 5 and leaf 12, respectively, compared to the WT (Fig. S7A). Similarly, *SAMBA* expression was diminished by 57.12% and 44.4% in *samba-3* for leaf 5 and leaf 12, respectively (Fig. S7B). However, the relative expression of *SAMBA* was also reduced by 47.7% in *samba-2* leaf 5, while the *samba-2* mutant did not reveal any growth defect (Fig. S7C). These data suggest that frame-shifts, as well as in-frame deletions, affect the stability of the transcript and that the downregulation of the *SAMBA* expression is not correlated with the severity of the observed phenotypes in the three *samba* alleles.

Next, we examined SAMBA protein formation by fusing the coding sequences without stop codon of the WT (positive control), *samba-1, samba-2* and *samba-3* alleles to a C-terminal YFP tag and overexpressing the constructs under the control of the CaMV 35S promoter in tobacco leaves. The samba-2 protein, which was expected to be about 0.96 kDa smaller than the WT because of the deletion of 10 AAs, showed a barely visible shift on Western blot (Fig. 5A). The samba-3 protein, which was expected to be about 6.31 kDa smaller than the WT and without the YFP tag because of the premature stop codon, would not be detected in the Western blot. The observed samba-3 protein (3,82 kDa smaller than the WT) could be explained when the protein started from the second methionine in SAMBA, of which the ATG, located at the start of exon 2, was recognized for translation re-initiation, resulting in an N-terminally truncated and C-terminally altered protein (Fig. 1C). Because only cDNA containing three exons was detected in WT and the *samba* alleles (Fig. S8), it was concluded that the truncated protein is not due to exon skipping but rather to translation re-initiation. In *samba-1*, the stretch of missense AAs reaches until the second methionine, most likely prohibiting the re-initiation of the translational machinery, explaining the absence of the N-terminally truncated SAMBA protein in *samba-1* on Western blot. Conversely, in *samba-3* the translational machinery is released from the mRNA after building in only 3 missense AAs, making the second methionine more accessible for translation re-initiation (Fig. 1C).

**Figure 5.**
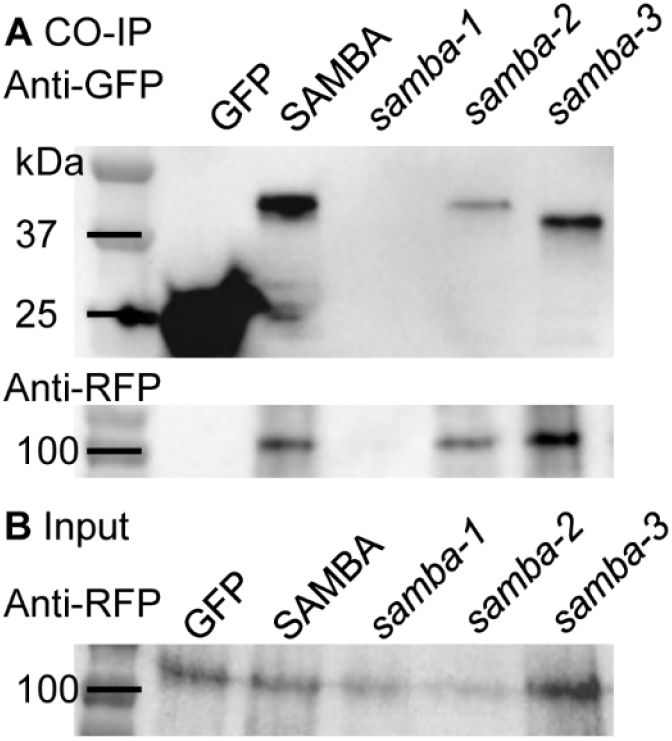
Co-immunoprecipitation results of C-terminally YFP-tagged WT SAMBA and *samba* alleles with C-terminally RFP-tagged APC3. Co-immunoprecipitation **(A)** and input of the extracted whole proteins **(B)**.

To analyze whether the mutated SAMBA proteins interact with the APC/C, a co-immunoprecipitation (Co-IP) of YFP tagged *SAMBA* (WT and the three alleles) with *APC3b* through its RFP tag were performed in tobacco leaves. Although APC3b-RFP proteins (107.05 kDa) were present in the input samples (Fig. 5B), they were detected after immunoprecipitation of both WT, *samba-2* and *samba-3*, indicating that the N-terminal and the C-terminal AAs of SAMBA were not essential for the protein interaction with the APC3b (Fig. 5A).

To further investigate the functionality of SAMBA in *samba-3*, we performed the SAMBA and *samba-3* AP/MS in maize callus using WT and the truncated SAMBA as it occurs in the *samba-3* mutant. Because of the one base deletion at second target site in *samba-3*, there is no stop codon at the end of *samba-3* coding sequence when the protein translation starts from the second methionine. The stop codon appears later in the 3’-UTR, causing an additional 15 AAs compared to the samba-3 (Fig. 1C). Therefore, we fused the full *samba-3* coding sequence and *samba-3* with 3’-UTR (*samba-3-UTR*) to the GS^rhino^ affinity tag next to the the SAMBA WT, and ectopically expressed these three constructs in maize callus for AP/MS. All APC/C members except the APC7 (did not found in samba-3) identified in the SAMBA WT were also retrieved both in samba-3 and samba-3-UTR, suggesting the mutated samba-3 or samba-3-UTR proteins still interact with the APC/C complex (Fig. S9, Supplemental Dataset 4). Moreover, 49 proteins were significantly enriched in the SAMBA WT compared to the *samba-3* and *samba-3-UTR* mutants including DRP1E, while 30 proteins were significantly enriched in the *samba-3* mutants including two UBIQUITIN6 proteins and a ubiquitin-specific protease. The altered enrichment in targets and UBIQUITIN6 suggests a distinct efficiency of the APC/C complex when bound to a mutated samba-3.

## Discussion

In Arabidopsis, SAMBA has been identified as a negative regulator of the APC/C, because its loss of function results in increased cell proliferation (Eloy et al., 2012). Here, we validated that the maize SAMBA ortholog also interacts with APC/C. Furthermore, *samba* loss of function in maize *samba-1* and *samba-3* mutants also accelerates cell proliferation during leaf development, indicating that, also in maize, SAMBA negatively regulates cell proliferation through its interaction with APC/C. However, despite this seemingly conserved role of SAMBA, the phenotypic read-out is distinct in Arabidopsis and maize mutants, which most likely results from several inter-species differences or a combination thereof. In maize, *samba* mutants show an increased cell production, due to a substantially faster cell cycle, resulting in more cells. Because the size of the division zone and expansion zone were reduced although not significantly, the cells did not elongate sufficiently and the final cell size was significantly smaller. In Arabidopsis, the increased cell division also resulted in smaller cells. For example, overexpression of the *E2F* transcription factors *(E2Fa (De Veylder et al., 2002)* and *E2Ff (Ramirez-Parra et al., 2004))* or *Cyclin D* (*CYCD*) (*CYCD3 (Menges et al., 2006) and CYCD4 (Kono et al., 2007)*) genes in Arabidopsis, stimulated cell division by driving the G1/S transition, and reduced or did not alter the cell DNA ploidy level, leading to reduced cell size. In Arabidopsis, where endoreduplication in leaves is much more pronounced than in some monocots such as maize, the cell size of the *samba* mutants increased through higher ploidy levels^21^. Besides endoreduplication-dependent developmental differences, also the timing at which the phenotypic effects upon *SAMBA* perturbation become apparent is different between maize and Arabidopsis. In Arabidopsis, the effects of *SAMBA* loss of function are most pronounced early in seedling development and decreased throughout further development^21^. In maize, however, *SAMBA* loss-of-function seedlings develop normally and show only a moderate reduction in final leaf length, while later in development more severe reductions of both leaf and internode length are observed. This might be explained by the *SAMBA* expression pattern differences between Arabidopsis and maize. In Arabidopsis, *SAMBA* shows high expression in all tissues of germinating seedlings but quickly diminishes in root tissues and becomes restricted to the hypocotyl at 8 days after stratification^21^. Conversely, in maize, *SAMBA* is more stably expressed throughout the entire development (Sekhon et al., 2011). The distinct developmental regulation of *SAMBA* indicates that in maize, *SAMBA* is not only instrumental for the cell divisions in organ primordia, but also during processes where cell division plays a role throughout development. The most conceivable phenotype upon *SAMBA* loss of function, being leaf and internode elongation and ligule formation, but also the less pronounced effects on leaf width, are associated with changes in the rate and/or orientation of cell divisions (Sylvester et al., 1990; McKim, 2019). Using affinity enrichment of the SAMBA-GS^rhino^ fusion protein from both embryogenic callus and leaf tissues, we were able to recover all APC/C subunits, including the CCS52 proteins. CCS52A proteins regulate the onset of the endocycle in Arabidopsis leaves and roots. Both loss-of-function mutants of *AtCCS52As* are smaller than WT, and their leaves contain more but smaller cells, whereas the double *ccs52a1/a2* mutant is lethal (Vinardell et al., 2003; Vanstraelen et al., 2009; Baloban et al., 2013). Downregulation of the rice *OsCCS52A* results in semi-dwarfism and smaller seeds with an endosperm defect in endoreduplication (Su’udi et al., 2012). The reduced plant height and leaf size are also observed in the rice *tillering and dwarf 1* (*tad1*) mutant, which is an ortholog of *CCS52A (Xu et al., 2012)*. The CCS52B protein is reported as a plant-specific APC/C activator (Tarayre et al., 2004) and the rice *ccs52b* mutant is semi-dwarf and shows narrow kernels (Su’udi et al., 2012). As the phenotype of the *ccs52* mutants in Arabidopsis and rice show similar growth defects as the maize *samba* mutant, it would be quite interesting to investigate the function of maize *CCS52* genes by generating the *ccs52* CRISPR knock-out mutants to further decipher the functions of maize APC/C. In addition, we identified several members of the DRP family, including DRP1A, DRP1E, DRP2A, DRP2B and DRP5B, as SAMBA interactors. All of these proteins, except DRP5B, have been shown to localize to the cell plate during cytokinesis (Park et al., 1997; Hong et al., 2003; Fujimoto et al., 2008) and mutations in *DRP1A* and *DRP1E* result in defective cell plate assembly and cytokinesis, as well as defects in cell expansion (Kang et al., 2003). Furthermore, CALLOSE SYNTHASE was identified as maize SAMBA interactor, which is involved in cell plate synthesis and is regulated by DRP proteins (Verma, 2001). Interestingly, APC3b, which has been shown to directly interact with SAMBA^21^, has also been found to be localized at the cell plate during the early telophase (Perez-Perez et al., 2008). In mammalian cells, DRP1 has been shown to be degraded by the APC/C in a D-box dependent way (Horn et al., 2011), and DRP2A, DRP2B, DRP5B and CALLOSE SYNTHASE also contain D-box motifs (Verma, 2001) and could, therefore, be direct targets of the APC/C. In *samba-3*, where the SHR1 domain was lacking, but (the majority of) the SHR2 domain was present, SAMBA was still able to interact with APC3, indicating that SHR2 is pivotal for SAMBA interaction with APC/C. Although the *samba-3* interacted with the APC/C, in total 79 proteins such as DRP1E and ubiquitin 6 were found significantly enriched in the SAMBA or *samba-3* callus, respectively, indicating the activity or efficiency of the APC/C were altered in *samba-3*. Performing AP/MS with bait proteins on callus tissue initiated from immature embryos during the early steps of maize transformation provided good insights into the interacting proteins and could help to circumvent the generation of transgenic plants (Dedecker et al., 2016).

Because of the phenotypic differences in timing and severity of seemingly similar *samba-1* and *samba-3* alleles, it became clear that addition or removal of one nucleotide at the same position not necessarily results in a “frameshift” mutant but can make a distinction between a true knock-out and the formation of a truncated, but partially functional protein. The observations that processes such as translation re-initiation and exon skipping might cause rather unexpected read-outs of genome editing were already made several times for animal tissues (Tang et al., 2018; Smits et al., 2019), but remained fragmentary in plants so far. Alternative splicing was shown to be the underlying reason why CRISPR/Cas9 induced frame shifts in the rice *OsIAA23* gene did not result in the expected severe phenotypes (Jiang et al., 2019). Here, we showed that besides alternative splicing, plants also can use translation re-initiation as a possibility to overcome CRISPR/Cas9 induced point-mutations. The understanding that in frame deletions as well as point mutations affect transcript stability and that plants possess of inherent strategies such as alternative splicing and translation re-initiation will create a novel awareness in the choice of gRNAs. Often gRNAs are designed close to the translational start site with the aim to create early frame-shifts and knock-out, but the *samba-3* allele showed that early mutations can cause translation re-initiation, which was in turn dependent on the number of missense AAs that shielded the re-initiation methionine. Taken together, our data show that caution should be taken in designing gRNAs and in interpreting the effects of mutations at the DNA level only.

## Materials and methods

### Growth conditions in growth chamber and greenhouse

Plants for leaf growth monitoring were grown under growth chamber conditions with controlled relative humidity (55%), temperature (24 °C day/18 °C night), and light intensity (170–200 μmol/m^2^/s photosynthetic active radiation at plant level) provided by a combination of high-pressure sodium vapor (RNP-T/LR/400W/S/230/E40; Radium) and metal halide lamps with quartz burners (HRI-BT/400W/D230/E40; Radium) in a 16-h/8-h (day/night) cycle. Plants for adult plant trait characterization were grown under controlled greenhouse conditions (26 °C/22 °C, 55% relative humidity, the light intensity of 180 μmol/m^2^/s photosynthetic active radiation, in a 16-h/8-h day/night cycle).

### Generation of *SAMBA* constructs

GRMZM2G157878 was identified as the maize SAMBA ortholog using PLAZA (Eloy et al., 2012; Proost et al., 2015). CRISPOR (Concordet and Haeussler, 2018) was used to determine possible gRNAs and their off-target scores in the coding sequence of *SAMBA*. Two gRNA sequences were chosen based on their location, MitSpecScore and off-target count: 35rev and 297fwd (Supplemental Dataset 5). These sequences target the coding sequence of *SAMBA* and do not overlap with other genes. The CRISPR construct containing these two gRNA sequences was generated according to Xing, H. L. et al (Xing et al., 2014), with minor modifications. Briefly, the kanamycin resistance cassette in the pBUN411 destination vector was replaced with a spectinomycin resistance cassette. This new vector is called pBUN411-Sp. Primers were designed that matched with the 35rev and 297fwd gRNA sequences, after which PCR was performed on the pCBC-MT1T2 plasmid, resulting in a fragment containing the desired target sites and the correct sites for ligation into the pBUN411-Sp destination vector using Golden Gate cloning (Invitrogen).

The coding sequences of *SAMBA, samba-3* and *samba-3-UTR* flanked by AttB1/AttB2 Gateway recombination sites were amplified from B104 WT cDNA using PCR, which was then ligated into pDONR221 entry vectors using Gateway BP reactions. The *SAMBA* entry vector together with the pUBI-L promoter (Coussens et al., 2012) and GS^rhino^ tag (Van Leene et al., 2015) were ligated into the pBb7m34GW destination vector (Karimi et al., 2013) to generate the pUBIL:*SAMBA*::GS^rhino^ (*SAMBA-GS*) expression vector by Multisite Gateway recombination (Invitrogen). This vector is codon-optimized for monocots and contains a Bar selection marker for the selection of transgenic plants using phosphinotricin (Karimi et al., 2013). Expression vectors for affinity enrichment experiments in callus tissues were generated by Multisite Gateway recombination (Invitrogen) into the in-house developed pBb-G7-BBMR-Ef1a destination vector. This vector contains a Bar selection marker but also contains a pEF1a:BBM cassette, allowing co-transformation of the morphogenic factor BABY BOOM, which can enhance callus growth.

### Maize transformation, genotyping and biomass generation

Immature embryos of the maize inbred line B104 were transformed by *Agrobacterium tumefaciens* cocultivation (Coussens et al., 2012). In short, immature B104 embryos were co-cultivated with *A. tumefaciens* for 3 days followed by 1-week growth on non-selective medium. Transformed embryogenic calli were subsequently selected on increasing concentrations of phosphinotricin. After shoot induction from the selected calli, transgenic T0 shoots were transferred to soil. At maturity, these T0 shoots were backcrossed with B104 WT, resulting in a collection of T1 seeds from multiple independent transgenic events. For the overexpression lines, this segregating population of 1:1 non-transgenic:transgenic plants were used for phenotyping and biomass generation for AP-MS. For the genome editing lines, the offspring containing the heterozygous mutant alleles that did not contain the T-DNA harboring CRISPR/Cas9 were self-crossed to identify WT plants and homozygous mutants. The segregating population was self-pollinated for phenotypic analysis in the next generation. Screening for mutations and genotyping of *samba* CRISPR mutants was performed by sequencing the genomic region containing both target sites after PCR amplification (Supplemental Dataset 5). To select suitable transgenic lines for AP-MS, the collection of T1 lines was first subjected to a segregation analysis to select single-locus insertion lines. Because the high accumulation of the fusion protein forms the basis of a successful affinity purification experiment, expression analysis was performed at the protein level using antibodies specific for the GS^rhino^ tags via western blotting. Based on this expression analysis, the transgenic lines showing the highest accumulation of the bait protein were selected to be upscaled for prototypical analysis and AP-MS. To perform AP-MS from division zone and expansion zone in triplicate, approximately 720 seeds were sown in small pots and grown in the growth chamber. When the first leaf was fully grown, a solution for the Bar selection marker, containing 1% w/v glufosinate ammonium (Sigma), was applied to its blade. After three days, non-transgenic plants showed severe first leaf necrosis while transgenic plants were resistant to this treatment, allowing easy genotyping of a large number of plants. Transgenic seedlings were harvested two days after leaf four appearance. The fourth leaf of each seedling was dissected and the division zone (first basal cm of the leaf) and the expansion zone (cm three to four) were cut out of the leaf growth zone. Harvested division and expansion zone samples were frozen in liquid nitrogen and pooled for all harvested plants. Harvested tissues were stored at -70°C.

The *SAMBA-GS5* line was crossed with *samba-1* and *samba-3*, and the heterozygote offspring was self-pollinated to generate homozygous WT plants and *samba-1* and *samba-3* mutants expressing the complementation construct.

The callus was generated by slight modifications to the transformation protocol (Coussens et al., 2012). Thirty immature embryos from three different pollinated ears were transformed. Embryos were cocultivated with *A. tumefaciens* for three days and subsequently transferred to a resting medium for 1 week. Transgenic calli were selected by transferring them to selection medium I for 2 weeks, after which the calli were transferred to selection medium II and incubated for two weeks. Non-transgenic callus tissue, which became necrotic due to phosphinothricin selection, was removed during each transfer step. Calli were then transferred to fresh selection medium II and incubated for one additional week, after which the healthy callus tissues were harvested, frozen in liquid nitrogen, ground to a fine powder (approximately 10g) using mortar and pestle and stored at -70°C.

### Affinity purification followed by mass spectrometry (AP-MS)

The AP-MS was performed by a slightly adjusted protocol (Nelissen et al., 2015). During protein extraction, Benzonase treatment was performed for 15 min instead of 30 min to shorten the duration of the purification protocol. The protein extracts were divided into three technical repeats, containing 65 mg protein extract each. To each technical repeat, 50 µl of home-made magnetic IgG beads, equilibrated in extraction buffer, were added and incubated for 45 min on a rotating device at 4°C. Beads were washed with 2x500 µl ice-cold extraction buffer and transferred to a fresh tube. Beads were washed again with 1x500 µl ice-cold extraction buffer and 1x500 µl ice-cold extraction buffer without detergents and final time with 800 µl 50 mM ammonium bicarbonate solution at pH 8. On bead digestion was performed in 50 µl of ammonium bicarbonate solution containing 4 µl of Trypsin/LysC (Promega) solution and incubated at 37°C for 3 h in a thermomixer at 1,000 RPM. After 3 h, the beads were separated from the digest and the digest was spiked with an additional 2 µl of Trypsin/LysC solution and left to incubate overnight at 37°C in a thermomixer at 1,000 RPM. Samples were dried in a SpeedVac system and stored for MS analysis.

### Liquid chromatography-tandem MS analysis

The digested peptide mixtures were injected into a liquid chromatography-tandem MS system using an Ultimate 3000 RSLCnano LC (Thermo Fisher Scientific) in-line connected to a Q Exactive (SAMBA callus versus GIF control group and SAMBA leaf) or Q Exactive HF Biopharma (SAMBA-3 mutants callus versus SAMBA WT) mass spectrometer (Thermo Fisher Scientific). The sample mixture was first bound on an in-house prepared trapping column (100 μm internal diameter (I.D.) × 20 mm, 5 μm beads C18 Reprosil-HD, Dr. Maisch). For Q Exactive, peptides were separated on an in-house prepared analytical column (75 μm I.D. × 150 mm, 5 μm beads C18 Reprosil-HD, Dr. Maisch) packed in the needle (PicoFrit SELF/P PicoTip emitter, PF360-75-15-N-5; New Objective). Peptides were separated through a linear gradient from 98% solvent A (0.1% formic acid in water) to 56% solvent B (0.1% formic acid in water/acetonitrile, 20/80 [v/v]) for 30 min at a flow rate of 300 nL/min. Next, the column was washed for 5min, hence reaching 99% solvent B. The operation mode of the mass spectrometer was set at the data-dependent, positive ionization mode, which automatically switches between MS and MS/MS acquisition for the 5 most abundant peaks in a given MS spectrum. The Q Exactive implemented source voltage was 2.8 kV and the temperature of the capillary was set at 275°C. After one MS1 scan (m/z 400 to 2000, AGC target 3 × 10^6^ ions, maximum ion injection time 80 ms) obtained at a resolution of 70,000 (at 200 m/z), up to 5 MS/MS scans (resolution 17,500 at 200 m/z) were obtained of the most intense ions that fitted the predefined selection criteria (AGC target 5 x 10^4^ ions, maximum ion injection time 60 ms, isolation window 2 Da, fixed first mass 140 m/z, spectrum data type: centroid, underfill ratio 2%, intensity threshold 1.3 × E4, exclusion of unassigned, 1, 5 to 8, >8 charged precursors, peptide match preferred, exclude isotopes on, dynamic exclusion time 12 s). The HCD collision energy was kept at 25% of the normalized collision energy. For Q Exactive HF Biopharma, the peptides were separated on a 250 mm Waters nanoEase M/Z HSS T3 Column, 100Å, 1.8 µm, 75 µm inner diameter (Waters Corporation) kept at a constant temperature of 50°C. Peptides were eluted by a non-linear gradient starting at 1% MS solvent B reaching 55% MS solvent B (0.1% FA in water/acetonitrile (2:8, v/v)) in 65 min, 97% MS solvent B in 70 minutes followed by a 5-minute wash at 97% MS solvent B and re-equilibration with MS solvent A (0.1% FA in water). The mass spectrometer was operated in data-dependent mode, automatically switching between MS and MS/MS acquisition for the 12 most abundant ion peaks per MS spectrum. Full-scan MS spectra (375-1500 m/z) were acquired at a resolution of 60,000 in the Orbitrap analyzer after accumulation to a target value of 3,000,000. The 12 most intense ions above a threshold value of 13,000 were isolated with a width of 1.5 m/z for fragmentation at a normalized collision energy of 30% after filling the trap at a target value of 100,000 for maximum 80 ms. MS/MS spectra (200-2000 m/z) were acquired at a resolution of 15,000 in the Orbitrap analyzer.

### Protein identification and label-free quantification

The raw files were processed using MaxQuant software (Tyanova et al., 2016a). The data was searched against the Zea mays B73 RefGen_v3 database (obtained from ftp://ftp.ensemblgenomes.org/pub/release-27/plants/fasta/zea_mays/pep/) using the built-in Andromeda search engine. A second database containing sequences of all possible types of contaminants that are often found with TAP or with proteomics experiments in general was added. These contaminants include a common repository of adventitious protein sequences, a list with proteins that are often found in proteomics experiments and are present as unavoidable contaminations of protein samples or by accident (The Global Proteome Machine, www.thegpm.org/crap/). In addition, tag sequences of many common tags and typical Tandem affinity purification (TAP) contaminants, including sequences of resin proteins or implemented proteases, were added. For the searches, only oxidation of methionines and N acetylation of protein N termini were accepted as variable modifications. A mass tolerance for precursor (peptide) ions was set at 4.5 ppm, whereas for fragment ions this was set at 20 ppm. The used enzyme was trypsin/P, permitting two missed cleavages as well as cleavage in cases where proline followed arginine or lysine. A target-decoy approach with a False Discovery Rate (FDR) of 1% was used to filter protein identifications and PSM. Label-free quantification was performed using the MaxLFQ algorithm integrated into MaxQuant (Cox et al., 2014).

Due to the high degree of protein sequence similarity between both CC52A1 and CC52A2 in Arabidopsis (86.4%) and the two maize CCS52A members (94.5%), it was impossible to determine which maize gene is the correct ortholog of which Arabidopsis gene, and to recognize the peptide in the AP/MS data. Therefore, GRMZM2G159427 was arbitrarily assigned as CCS52A1 and GRMZM2G064005 as CCS52A2, and both genes were identified as CCS52As in the AP/MS data.

### Statistical analysis of label-free quantification

The identification of significantly enriched proteins in the samples was performed using Perseus (Tyanova et al., 2016b). The MaxQuant output files were loaded into Perseus and LFQ intensities were log2 transformed, after which samples were grouped in bait and control, or in division zone and expansion zone. Proteins that did not contain at least two valid values in at least one group were filtered out and missing LFQ values were imputed from the normal distribution, as previously described (Smaczniak et al., 2012; Wendrich et al., 2017). A Student’s *t*-test was performed to determine significantly enriched proteins in either developmental zone, or a permutation-based FDR correction (FDR=0.01, S0=1) was applied to correct for multiple hypothesis testing. Significantly enriched proteins after affinity enrichment of SAMBA-GS^rhino^ in callus were determined using a control dataset from the non-SAMBA-related proteins GIF2-GS^rhino^ and GIF3-GS^rhino^ AP/MS transformed embryogenic callus as a negative control. Additionally, non-specific and sticky proteins were marked in the lists, based on Supplemental Dataset 3, which is assembled from a set of control affinity purification experiments from maize leaf tissues (Besbrugge et al., 2018) and a set of proteins whose Arabidopsis orthologs were included in a previously published background list based on 543 TAP purifications over 155 different bait proteins (Van Leene et al., 2015).

### Quantitative RT-PCR

Expression analysis of *SAMBA* was checked by qRT-PCR. Total RNA was extracted from each repeat using Trizol (Life Technologies, Invitrogen) and DNA was removed by RQ1 DNase (Promega) treatment. Preparation of cDNA was performed using the iScript cDNA synthesis kit (BioRad) according to manufacturer’s recommendations starting with 1µg of RNA. qRT-PCR was performed with the LightCycler 480 Real-Time SYBR Green PCR System (Roche), used primers are listed in (Supplemental Dataset 5). For qPCR on maize samples, we used *18S RNA* as the housekeeping gene.

### Flow Cytometry

For flow cytometry analysis, at least three leaves or stem samples per time point of a biological repeat (n = 3) were chopped with a razor blade in 200 ml of Cystatin UV Precise P Nuclei Extraction buffer (Partec), followed by the addition of 800 ml of staining buffer and filtering through a 50-mm filter. Nuclei were analyzed with the Cyflow MB flow cytometer (Partec) and the FloMax software.

### Anatomical analysis

The 2^nd^ cm of the *samba-1* mature ninth leaf blade was cut into cross slides by hand section. Segments were soaked in 100% ethanol for discoloration and then transferred into the ACA staining solution (0.5% w/v Astra blue, 0.5% w/v Chrysopidae and 0.5% w/v acridine red solution mixed at a volume ratio of 16:1:1) for 5 min. The stained sections were washed 3 times, 5 min per time in water and mounted in 50% glycerol solution for imaging.

### Co-IP assays

The coding sequence without stop codon of WT *SAMBA, samba-1, samba-2, samba-3* and were fused with a 3’-YFP tag; the *APC3b* was fused with a 3’-RFP tag. All genes were cloned behind the 35S promoter in the pBb7m34GW destination vector (Karimi et al., 2013). All these constructs, including the empty p35S::YFP vector were introduced into *A. tumefaciens* cells. All transient expression assays were performed as described previously (Marin-de la Rosa et al., 2015). Tobacco leaves were harvested 48 h after infiltration and were ground in liquid nitrogen, homogenized in the extraction buffer (50 mM Tris·Cl pH 7.5, 1 mM EDTA, 75 mM NaCl, 0.1% Triton X-100, and protease inhibitor cocktail (Sigma)). The supernatant was incubated with the GFP-Trap agarose beads (Chromotek) for 2 h at 4 °C. After washing, beads were eluted by boiling with 2× SDS sample buffer for 10 min. Samples were separated on 4–20% Mini-PROTEAN® TGX Stain-Free(tm) Protein Gels (Bio-Rad) and then immunoblotted with anti-GFP and anti-RFP antibody (Abcam, 1:2000 dilution).

## Acknowledgements

The research was funded by the European Research Council under the European Community’s Seventh Framework Programme [FP7/2007-2013] under ERC grant agreement n° [339341-AMAIZE]11 and by the ‘Bijzonder Onderzoeksfonds Methusalem Project’ (no. BOFMET2015000201) of Ghent University. Pan Gong received a scholarship from the China Scholarship Council (CSC no. 201606320217). The authors thank Dr. Bert De Rybel, Helena Arents and Ji Li for helping with the leaf hand section and ACA staining, Dr. Hannes Vanhaeren, Dr. Ting Li, Ying Chen and Reinout Laureyns for helping with the Co-IP Assay, Rudy Vanderhaeghen for callus generation, Thomas Eeckhout for sharing the pBUN411-Sp vector and VIB Proteomics core facility for analyzing the maize AP-MS samples.

## Author contributions

P.G., M.B., L.P., G.D.J., D.I. and H.N. conceived and planned the experiments. AP.G., M.B., K.D., J.D.B., G.P., S.A, G.C. carried out the experiments. P.G., M.B., K.G., D.E., M.V.L., G.D.J., D.I and H.N. contributed to the interpretation of the results. P.G., M.B., D.I. and H.N. took the lead in writing the manuscript. All authors provided critical feedback and helped shape the research, analysis and manuscript.

## Figure legends

**Figure S1. Expression and phenotypic analysis of the *SAMBA-GS* transgenic overexpression lines. (A)** Western blotting. The molecular weight of the fusion proteins is 32 kDa. The first centimeter of 10-days-old shoots was harvested for protein extraction. Proteins were detected using peroxidase anti-peroxidase soluble complex antibody specific for the GS^rhino^-tag. **(B)** Final leaf 4 length of *SAMBA-GS5*. **(C)** Plant height (from soil base to highest leaf collar) of *SAMBA-GS5* at silking stage. Error bars represent standard error, n ≥ 3.

**Figure S2. Label-free quantification of APC/C components and regulatory proteins co-purified with SAMBA affinity enrichment using AP-MS on the division (DZ) and expansion zone (EZ) of the growing maize leaf**. The difference in average log2 LFQ intensities between DZ and EZ is plotted versus the significance (threshold FDR = 0.01 and S0 = 1; Supplemental dataset 2). The curve indicates the permutation-based FDR threshold. Positions of SAMBA, APC and CCS52 family proteins are marked in blue on the plot.

**Figure S3. Average internode and leaf mature epidermal cell length of *samba-1* and *samba-3* mutants**. The average cell length of *samba-1* internode 9 (A), *samba-3* internode 12 (B), *samba-1* leaf 12 (C) and samba-3 leaf 12 (D). The central three centimeters of well-developed internodes and leaves’ epidermal cell length were matured. Error bars represent standard error and significant differences were determined using Student’s *t*-test: *, P<0.05, **, P<0.01 (n=3).

**Figure S4. Representative root systems of *samba-1* (A), *samba-3* (B) and *samba-1/3* (C) mature plants at tasseling stage of the respective WT**. Bar =5 cm

**Figure S5. Microscopic comparison of mature leaves and internodes of WT and *samba-1* mutants**. Cellular imprint of mature internode 9 epidermis of WT **(A)** and *samba-1* **(B)**. Cellular imprint of mature leaf 9 epidermis of WT **(C)** and *samba-1* **(D)**. Macrohairs (MH) and bicellular microhairs (BH) on WT **(E)** mature leaf blade and prickle hairs (PH) on *samba-1* mutant leaf blades **(F)**. The asterisk indicates the location of bulliform cell files. Macrohairs and prickle hairs on WT **(G)** and *samba-1* **(H)** leaf margins. Leaf 9 midrib section and ACA staining of WT **(I)** and *samba-1* **(J)**.

**Figure S6. Ploidy distribution of the mature leaf and internode cells in *samba-1* and *samba-3* mutants determined by flow cytometry**. The cell ploidy distribution of *samba-1* leaf 12 (A), *samba-3* leaf 12 (B), *samba-1* internode 9 (C) and samba-3 internode 12 (D). Error bars represent standard error and significant differences were determined using Student’s *t*-test: *, P<0.05, **, P<0.01 (n=3).

**Figure S7. The relative expression level of *SAMBA* determined by Q-RT-PCR in *samba-1* (A), *samba-3* (B) and *samba-2* (C) relative to the respective segregating WT plants**. Error bars represent standard error and significant differences were determined using Student’s *t*-test: *, P<0.05 (n = 3).

**Figure S8. The PCR for the UTRs of *SAMBA* in WT plants and the *samba* alleles on cDNA from the leaf 4 division zone. (A)** Agarose gel electrophoresis (1%) of PCR products. **(B)** Sequencing results of the PCR.

**Figure S9. Label-free quantification of APC/C components and regulatory proteins co-purified with SAMBA, samba-3 and samba-3-UTR affinity enrichment using AP-MS on the maize callus**. The differences in average log2 LFQ intensities between WT SAMBA and samba-3 **(A)**, WT and samba-3-UTR **(B)** are plotted versus the significance (threshold FDR = 0.02 and S0 = 2; Supplemental dataset 4). The curve indicates the permutation-based FDR threshold. Positions of APC/C and CCS52 family proteins are marked in red on the plot.

**Supplemental dataset 1: Protein identifications through affinity enrichment from maize callus by using SAMBA-GSrhino as bait protein**.

**Supplemental dataset 2: Protein identifications through affinity enrichment from leaf division and expansion zone using SAMBA-GSrhino as bait protein**.

**Supplemental dataset 3: Lists of non-specific and sticky proteins used for background filtering**.

**Supplemental dataset 4: Protein identifications through affinity enrichment from**

***SAMBA* and *samba-3* mutants callus using SAMBA-GSrhino as bait protein**.

**Supplemental dataset 5: Nucleotide sequences of the primers used for cloning and qPCR**.

